# Unveiling the co-phylogeny signal between plunderfish *Harpagifer* spp. and their gut microbiomes across the Southern Ocean

**DOI:** 10.1101/2023.04.18.537398

**Authors:** Guillaume Schwob, Léa Cabrol, Thomas Saucède, Karin Gérard, Elie Poulin, Julieta Orlando

## Abstract

Understanding the factors that sculpt fish gut microbiome is challenging, especially in natural populations characterized by high environmental and host genomic complexity. Yet, closely related hosts are valuable models for deciphering the contribution of host evolutionary history to microbiome assembly, through the underscoring of phylosymbiosis and co-phylogeny patterns. Here, we hypothesized that the recent allopatric speciation of *Harpagifer* across the Southern Ocean (1.2–0.8 Myr) will promote the detection of robust phylogenetic congruence between the host and its microbiome.

We characterized the gut mucosa microbiome of 77 individuals from four field-collected species of the plunderfish *Harpagifer* (Teleostei, Notothenioidei), distributed across three biogeographic regions of the Southern Ocean. We found that seawater physicochemical properties, host phylogeny and geography collectively explained 35% of the variation in bacterial community composition in *Harpagifer* gut mucosa. The core microbiome of *Harpagifer* spp. gut mucosa was characterized by a low diversity, mostly driven by selective processes, and dominated by a single *Aliivibrio* taxon detected in more than 80% of the individuals. Almost half of the core microbiome taxa, including *Aliivibrio*, harbored co-phylogeny signal at microdiversity resolution with *Harpagifer* phylogeny. This suggests an intimate symbiotic relationship and a shared evolutionary history with *Harpagifer*.

The robust phylosymbiosis signal emphasizes the relevance of the *Harpagifer* model to understanding the contribution of fish evolutionary history to the gut microbiome assembly. We propose that the recent allopatric speciation of *Harpagifer* across the Southern Ocean may have generated the diversification of *Aliivibrio* into patterns recapitulating the host phylogeny.

**Importance:** Although challenging to detect in wild populations, phylogenetic congruence between marine fish and its microbiome is critical, as it allows highlighting potential intimate associations between the hosts and ecologically relevant microbial symbionts.

Through a natural system consisting of closely related fish species of the Southern Ocean, our study provides foundational information about the contribution of host evolutionary trajectory on gut microbiome assembly, that represents an important yet underappreciated driver of the global marine fish holobiont. Notably, we unveiled striking evidence of co-diversification between *Harpagifer* and its microbiome, demonstrating both phylosymbiosis of gut bacterial communities, and co-phylogeny of specific bacterial symbionts, in patterns that mirror the host diversification. Considering the increasing threats that fish species are facing in the Southern Ocean, understanding how the host evolutionary history could drive its microbial symbiont diversification represents a major challenge to better predict the consequences of environmental disturbances on microbiome and host fitness.

## 1. Background

The implication of the microbiome in facilitating or responding to host evolutionary processes is one of the burning points of holobiont studies (1–3). Under the holobiont concept, the reciprocal evolution of host and microbiome genomes, namely co-evolution, is associated with several key life history traits such as obligate symbiosis, vertical inheritance, metabolism cooperation, reproduction control and co-diversification (4, 5). Co-diversification, defined as the parallel and concomitant diversification of the host and symbiont lineages through a history of constant association (6), constitutes the most investigated trait to get evidence of potential co-evolution in natural macroorganisms’ populations (7–10). Yet, directly testing for co-diversification between a host and its microbiome remains challenging, because signal might be absent or weak, and available data from natural populations are usually limited (11). Thus, two empirical patterns are classically recognized, coined phylosymbiosis and co-phylogeny, to evidence the impact of the evolutionary interactions on holobiont assemblage (11–13). Phylosymbiosis refers to a congruence pattern between the phylogeny of host species and the clustering of microbial community structure, observed at one moment in time and space (14, 15). However, this pattern does not imply any stable evolutionary association between a host and its microbiome along time (16). Practically, phylosymbiosis is reflected by higher similarity of microbial communities within the same host species than between different host species, and by genetic differences among hosts that are consistent with the compositional differences in their microbiomes (17). By contrast, co-phylogeny involves parallel evolutionary history of host species and specific microbial symbionts (*i.e.* microbial lineages conforming the microbiome) (18). A co-phylogenetic signal is elucidated by congruent topologies of host species and specific symbionts phylogenies, by which interacting partners shared similar positions in their respective trees (11). The screening for phylosymbiosis and co-phylogeny signals in complex and uncharacterized holobionts has led to the identification of specific microbes, with potentially highly relevant ecological role in the host (10, 19–21).

Detecting robust phylosymbiosis and co-phylogeny signals in wild species populations is challenging, due to the complexity of natural holobiont systems related to uncontrolled sources of microbial variability (15, 22). To tackle this issue, several approaches have been proposed, such as focusing on the core microbiome (*i.e.* common microbial taxa across diverse environments) (22, 23), and on the mucosa resident microbiome (*i.e.* autochthonous and supposedly temporally stable in the intestinal mucus) (17, 24, 25). These host-specific sub-communities, less impacted by external environment and diet (26–28), likely fulfill critical functions for the host and contribute to its fitness and evolution (5, 17, 29), and hence are more susceptible to present co-phylogeny patterns (30). Moreover, in anciently diverged hosts species with long-term evolution from the last common ancestor, the phylosymbiosis signal can be blurred by evolutionary history events (*e.g.* host-swap, symbiont extinction or phylogenetically non-congruent symbiont speciation events) (15, 31). Several authors advocated working with recently diverged, and thus genetically closely-related species, characterized by stronger phylosymbiosis pattern (32), to better delineate the effects of evolutionary history and ecology of the host on microbiomes assembly (3, 22, 33, 34).

The fish fauna of the Southern Ocean (SO) is dominated by the perciform suborder Notothenioidei (teleost fishes), which constitute 90% of the fish biomass and up to 77% of the species diversity of the Antarctic continental shelf (35, 36). To date, the microbial communities associated to the notothenioids have received very little attention, and has been mainly characterized through cultural-dependent methods (37–39). The gut microbiome of only two species from West Antarctic Peninsula, *Notothenia coriiceps* and *Chaenocephalus aceratus,* has been analyzed through 16S rRNA gene-based clone libraries, revealing a low poor microbial richness and a strong dominance of the family *Vibrionaceae* represented by *Vibrio* and *Photobacterium* genera (35). However, none of these studies addressed co-diversification within the notothenioid holobiont.

Within the Notothenioidei suborder, the monogeneric family of the *Harpagiferidae* groups 12 nominal species, each one distributed in a specific region of the SO, such as *Harpagifer antarcticus* in Antarctica (40), *H. geogianus* in the South Georgia Islands (41), *H. bispinis* in Patagonia (42), and *H. kerguelensis* in Kerguelen Islands (43). The diversification of the *Harpagifer* genus occurred 1.2–0.8 Myr ago, from Antarctica towards the Patagonia and sub-Antarctic areas during the Pleistocene (44, 45). This recent divergence makes *Harpagifer* an interesting model to explore co-diversification hypothesis. Despite contrasting environmental conditions due to their distinct geographical distribution, these closely related species have the same trophic positioning, and live in intertidal and shallow subtidal habitats (46, 47). Thus, we expect to detect shared microbial taxa, conforming the core of the gut mucosa microbiome (‘GMM’ hereafter) among *Harpagifer* species. The slow evolution of the 16S rRNA gene marker of bacteria, at 1-2% per 50 Myr on average (48), precludes exploring pattern of co-phylogeny among symbionts and their host species that have diverged in shorter timescales when considering the classical 97% similarity bacterial OTUs (49). Thus, the co-phylogenetic signal of *Harpagifer* species and their microbiomes will be explored at a microdiversity level (50). By characterizing through 16S rRNA gene sequencing the GMM of wild-caught individuals from four species of *Harpagifer*, we aim (i) to evaluate the contribution of host identity and phylogeny on gut microbiome composition compared to the environment and the geography, and (ii) to test the hypothesis of a co-phylogenetic signal between *Harpagifer* species and shared members of their gut microbiomes (*i.e.* core microbial taxa).

## 2. Methods

### 2.1 *Harpagifer* spp. individuals sampling and dissection

Individuals of the fish species *H. bispinis*, *H. georgianus*, *H. kerguelensis* and *H. antarcticus* were sampled between 2015 and 2021 from 12 localities of the SO, including two localities in the Chilean Patagonia (PAT1 and PAT2), five localities in South Georgia, one locality in the Kerguelen Islands (KER) and four localities in the West Antarctic Peninsula (WAP1-WAP4), respectively (Figure 1, Table 1). Individuals were euthanized using buffered seawater containing >250 mg/L benzocaine (BZ-20^®^, Veterquimica^®^) and were then conserved in absolute ethanol at 4°C until dissection. Once at the lab, the *Harpagifer* individuals were aseptically dissected to remove the intestinal content, and the gut mucosa were gently rinsed with nuclease-free sterile water (Winkler) and stored at -20°C until DNA extraction.

**Figure 1.**
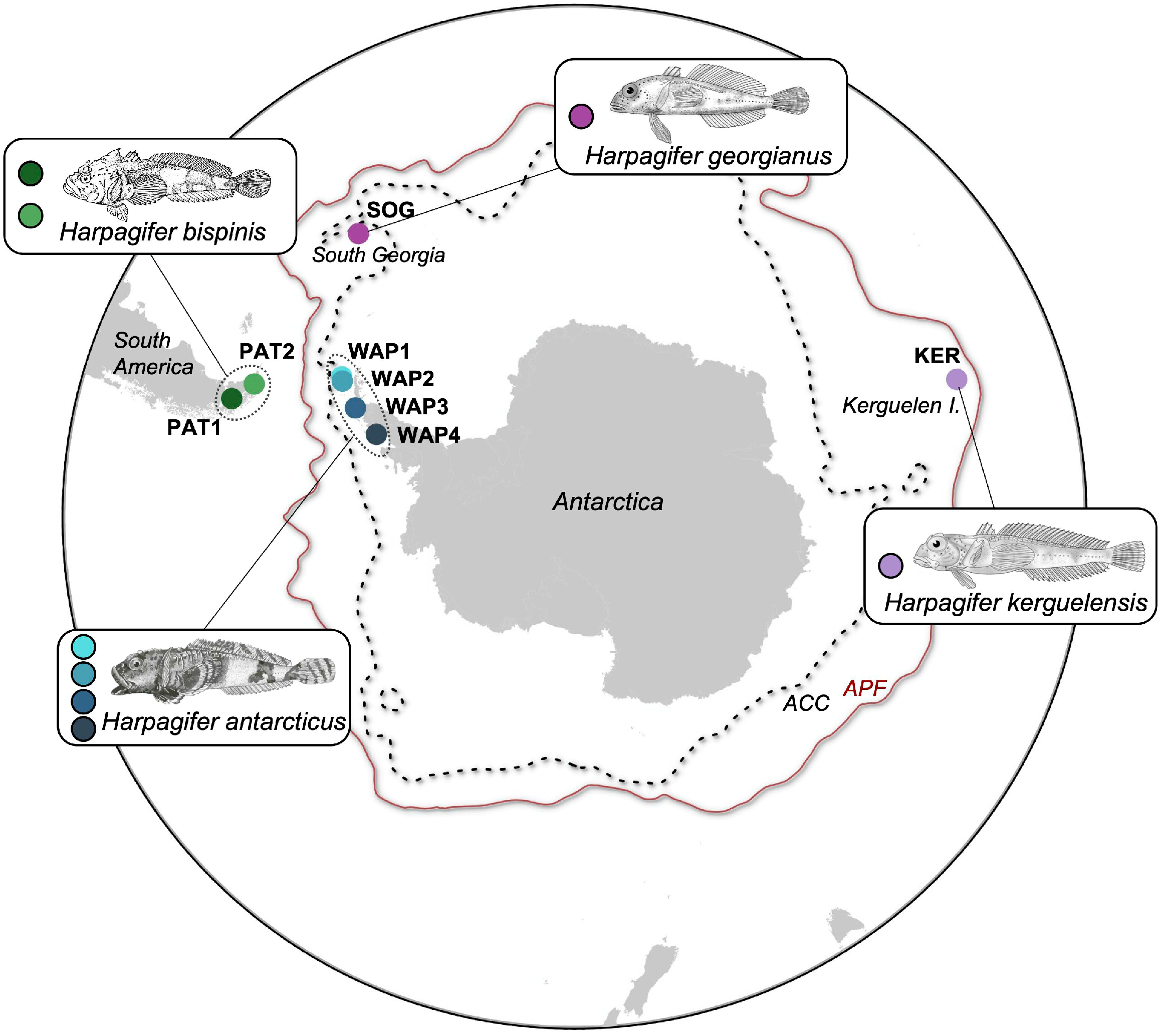
Sampled *Harpagifer* sp. populations across the Southern Ocean.

**Table 1.** Sampling locations of Harpagifer species and overview of amplicon sequencing data.

| Species | Region | Locality | Date | GPS coordinates | N | N seq. (Relat. Abund.) |
| --- | --- | --- | --- | --- | --- | --- |
| *Harpagifer antarcticus* | West Antarctic Peninsula (**WAP**) | Antarctic China Base "Great Wall", Fildes Peninsula, King George Island, South Shetland Islands (**WAP1**) | 01-2020 | 62°12'28.20"S<br>58°57'38.00"W | 10 | 318876 (10.7) |
|  |  | Antarctic Chilean Base "Captain Arturo Prat", Iquique Cove, Greenwich Island, South Shetland Islands (**WAP2**) | 01-2018 | 62°28'43.84"S<br>59°40'14.90"W | 10 | 490478 (16.4) |
|  |  | Antarctic Chilean Base "Yelcho", South Bay, Doumer Island, Antarctic Peninsula (**WAP3**) | 01-2018 | 64°52'33.13"S<br>63°35'00.96"W | 8 | 373151 (12.5) |
|  |  | Horseshoe, Marguerite Bay, Antarctic Peninsula (**WAP4**) | 01-2022 | 67°53'32.82S<br>67°24'17.34"W | 10 | 247211 (8.3) |
| *Harpagifer kerguelensis* | Kerguelen Islands (**KER**) | Port-aux-Français, Gulf of Morbihan, French Southern and Antarctic Lands | 12-2015 | 49°21'13.32"S<br>70°13'56.76"E | 11 | 255340 (8.5) |
| *Harpagifer georgianus* | South Georgia Island (**SOG**)* | Esbensen Bay | 11-2021 | 54°52'0.96"S<br>35°57'46.98"W | 7 | 297991 (10.0) |
|  |  | Gold Harbour | 11-2021 | 54°36'59.16"S<br>35°55'15.899"W |  |  |
|  |  | Albatross Island, Sunset Fjord | 11-2021 | 54°1'37.56"S<br>37°19'26.399"W |  |  |
|  |  | Sector Godthul | 11-2021 | 54°17'24.12"S<br>36°17'6.299"W |  |  |
|  |  | Luck Point, Bay of Isles | 11-2021 | 54°3'37.92"S<br>37°15'54.9"W |  |  |
| *Harpagifer bispinis* | Chilean Patagonia (**PAT**) | Sector "Aguas Frescas", Punta Arenas, Strait of Magellan, Magallanes and Chilean Antarctica Region (**PAT1**) | 02-2021 | 53°26'00.23"S<br>70°58'24.37"W | 11 | 558754 (18.6) |
|  |  | Sector "Corrales Viejos", Puerto Williams, Navarino Island, Magallanes and Chilean Antarctica Region (**PAT2**) | 04-2021 | 54°55'56.89"S<br>67°28'20.89"W | 10 | 448484 (15.0) |

N: number of individuals per species, Nseqs: total number of sequences after bioinformatic processing, Relat.
Abund.: relative abundance in the whole dataset, \*: Given the low number of individuals per locality within the South Georgia Island, the samples from the 5 localities were pooled.

### 2.2 Genomic DNA extraction, PCR amplification and amplicon-sequencing analysis

DNA was extracted from gut mucosa samples using the DNeasy^®^ PowerSoil^®^ Pro Kit (Qiagen), with a preliminary incubation at 65°C for 10 min followed by a homogenization step using a FastPrep-24™ bead beating grinder (MP Biomedicals™). The V3-V4 region of the bacterial 16S rRNA gene was amplified by touchdown PCR using the modified Bakt_341F/Bakt_805R primer pair (51). PCR products were purified and sequenced using the paired-end sequencing technology (2x300 bp) on the Illumina MiSeq Sequencer at the University of Wisconsin – Madison Biotechnology Center’s DNA Sequencing Facility (USA). Reads of 16S rRNA gene were processed through MOTHUR (v1.48.0) (52), using the trimming criteria detailed in Schwob et al. (2019) (53). Processed sequences were clustered into OTUs at 97% similarity threshold similarity, discarding the OTUs conformed by a single sequence.

The host mitochondrial COI gene was amplified from the same DNA samples, using the FISH-F2/HCO2198-R primer pair (54, 55). Amplicons were purified and sequenced in both directions at Macrogen Inc. (South Korea) using Sanger technology. Sequences of COI gene from the *Harpagifer* individuals were aligned and polymorphic sites were visually checked in PROSEQ (56).

### 2.3 Host genetic diversity, structure and phylogeny reconstruction

The classification of the edited COI sequences of *Harpagifer* individuals into haplotypes was performed using ARLEQUIN (v3.5.2). The haplotype network was reconstructed using the Median Joining method with the software Populational Analysis with Reticulate Trees (PopART, v1.7.0) (57). The phylogenetic tree was reconstructed using PhyML (58), and rooted using the midpoint rooting method (59) to avoid long-branch attraction.

Pairwise genetic distances among *Harpagifer* individuals were computed from the reconstructed host phylogenetic tree with the APE package (v5.6-2) in R (v4.1.2).

### 2.4 Statistic analyses for phylosymbiosis detection

The bacterial OTU table was rarefied at 5,750 sequences and converted into Bray-Curtis dissimilarity distances. To test the effect of host species identity on *Harpagifer* GMM composition, a permutational multivariate analysis of variance (PERMANOVA) was performed using VEGAN (v2.6-2) and PAIRWISEADONIS (v0.4) R packages, respectively. A set of 14 environmental abiotic variables presumed important in influencing microbiome structure, was extracted for each sampling locality from the Bio-ORACLE database (60) (Supplementary Figure 1). All the environmental variables, standardized to a mean of zero and a standard deviation of one, were analyzed using a principal component analysis (PCA). The scores of the samples on the first two principal components (capturing 91% the variability), were transformed into Euclidean distance using the *vegdist* function of the R package VEGAN (v2.6-2) and used as environmental distance matrix (Supplementary Figure 1). The geographic distances among sampling localities were obtained by converting longitude and latitude coordinates into kilometers with the *earth.dist* function implemented in the FOSSIL package (61), followed by a transformation with the Hellinger method using the *decostand* function of the VEGAN package in R. All matrices were standardized using the *scale* function implemented in R. The correlation between dissimilarity distances of the *Harpagifer* GMM with each one of the three explicative distance matrices (*i.e.* environmental, geographic and host phylogeny) was examined with Mantel and partial Mantel tests implemented in VEGAN, using Pearson correlation and 9999 permutations (62). Further, the respective contribution to GMM composition of these explanatory matrices was inferred using the distance-based Multiple Matrix Regression with Randomization (MMRR) approach (63), implemented in the R package POPGENREPORT (v3.0.7).

### 2.5 Core microbiome definition and neutral model fitting

To identify bacterial taxa that are common to the eight *Harpagifer* populations, a core microbiome was defined at the OTU level, based on a minimum prevalence criterion >40% across all gut mucosa samples from our dataset (23). The relationship between the prevalence and abundance of core microbiome OTUs was compared to the neutral community model proposed by Sloan et al. (64) and formalized in R by Burns et al. (65). Well-predicted OTUs (*i.e.* with abundance and prevalence comprised within the 95% confidence limits of the model) are supposed to be driven by stochastic factors (*e.g.* dispersal and ecological drift), while OTUs that deviated from the 95% confidence interval of the neutral model predictions are more likely influenced by deterministic factors (*e.g.* host selection).

### 2.6 Microdiversity analysis and co-phylogeny testing

The microdiversity of each core OTU (n=17) was resolved into oligotypes through the Minimum Entropy Decomposition (MED) algorithm developed by Eren et al. (50). Briefly, MED allows discriminate the biologically meaningful microdiversity contained within one OTU from the stochastic noise caused by random sequencing errors (66). The oligotypes’ phylogenetic tree of each core OTU from the core microbiome was inferred using PhyML 3.0 (67). Co-phylogeny patterns between the bacterial oligotypes and *Harpagifer* species phylogenetic trees were tested through a total of 10 runs with 999 permutations of the *Parafit* function, implemented in the APE package (v5.6-2). The *Parafit* function returned the relative contribution of each individual host-oligotype link to the co-phylogenetic model, with their associated *p*-values adjusted using the Benjamini-Hochberg procedure. For the graphical representation of the co-phylogeny pattern for the most abundance core OTU (OTU2), links with *p*-values > 0.055 were pruned and the tanglegram was edited using the R package PHYTOOLS (v1.0-3) (68). To further investigate the mechanism underlying the co-phylogenetic pattern of OTU2, we used the Procrustean Approach to Co-phylogeny (PACo) (69) implemented in R (70) to test the dependence of one phylogeny on the other. The best model was determined by comparing the phylogenetic congruences obtained through 20,000 permutations using the pairwise permutation test implemented in the R package RCOMPANION (v2.4.16). The haplotype network of the OTU2 was reconstructed as previously described for the host (section 2.3). Distance-based redundancy analysis (db-RDA) implemented in the R package VEGAN was used to quantify the contribution of the *Harpagifer* species to the variations in OTU2 oligotypes composition.

## 3. Results

### 3.1 *Harpagifer* genetic structure

In total, 81 sequences of *Harpagifer* COI were obtained, defining 25 haplotypes that conformed three distinct haplogroups corresponding to Patagonian populations (PAT1 and PAT2, hereafter called PAT), Kerguelen Islands population (KER), and Antarctic and South Georgia populations (WAP and SOG) (Supplementary Figure 2 and Supplementary Table 1).

### 3.2 *Harpagifer* species identity influences its gut mucosa microbiome

A total of 77 gut mucosa samples were successfully processed through metabarcoding, resulting in 2,990,285 cleaned sequences partitioned into 34,419 OTUs at 97% (Table 1). A weak but significant effect of *Harpagifer* species identity on GMM composition was detected (PERMANOVA, F-statistics=4.46, R^2^=0.15, *p*<0.001). All pairwise comparisons of GMM compositions among *Harpagifer* species were significantly different (pairwise PERMANOVA, *p*<0.04) except between *H. antarcticus* (WAP) and *H*. *georgianus* (SOG) (Supplementary Table 2), mirroring the absence of genetic differentiation among host haplotypes from these two regions (Supplementary Figure 2). The differences in microbiome composition between gut mucosa from KER and PAT were weak (pairwise PERMANOVA, *p*=0.04), echoing the relatively more similar seawater properties of these two regions (Supplementary Figure 1).

### 3.3 Robust phylosymbiosis of *Harpagifer* species and their gut mucosa microbiome

To further predict the assembly mechanisms of *Harpagifer* GMM, we tested whether the Bray-Curtis dissimilarity distance correlates with the geographic, environmental and host phylogenetic distance matrices. The Mantel correlations tests revealed that environment, host phylogeny and geography explained significant and relatively comparable amounts of variability in GMM of *Harpagifer* species (Table 2). When evaluating the independent effect of each explanatory matrix by controlling for the influence of other matrices variations with partial Mantel tests, both the host phylogeny and the environment (and to a lower extent the geography) still significantly correlated with beta-diversity variations of *Harpagifer* GMM, even though with reduced explanatory power (Table 2). When examined through the MMRR analysis, the relative contribution of each matrix on bacterial composition of *Harpagifer* GMM remained within the same order of magnitude as previously obtained with Mantel tests, with the highest contribution for the environment (coefficient of 0.14, *p*<0.001), followed by the host phylogeny (coefficient of 0.10, *p*<0.001) and geography (coefficient of 0.08, *p*<0.001). The Mantel test performed with the combined distance matrix computed from the three original ones weighted by their respective MMRR coefficient provided the best-fit model (R^2^=0.35, p<0.001) (Figure 2).

**Figure 2.**
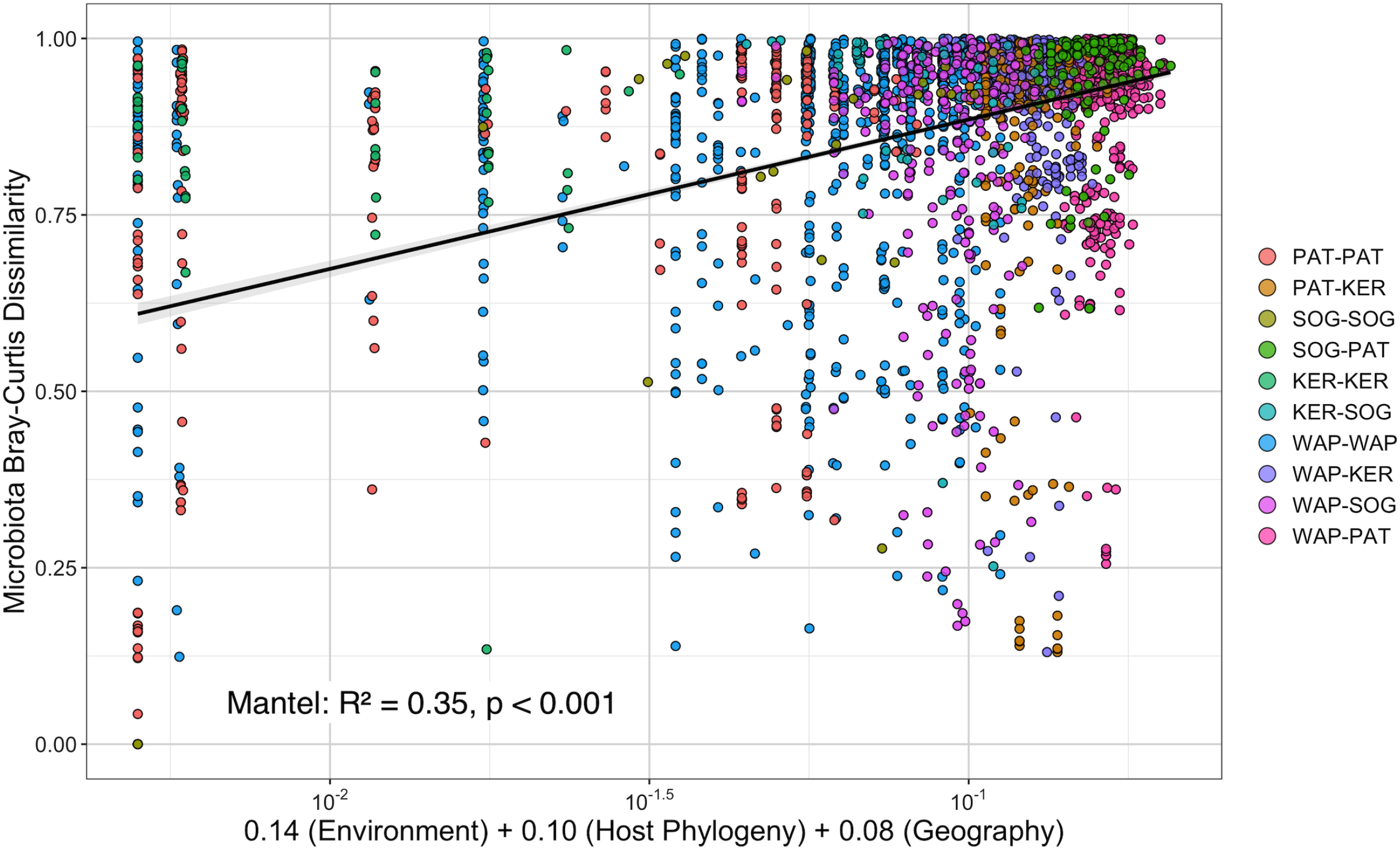
Scatter plots showing the relationship between the Bray-Curtis dissimilarity of *Harpagifer* mucosa microbiome and the joint effect of host phylogenetic, geographic and marine environmental distances based on the results of a Multiple Matrix Regression with Randomization analysis (MMRR). Color of the points represents pairwise comparisons among sampling localities.

**Table 2.**
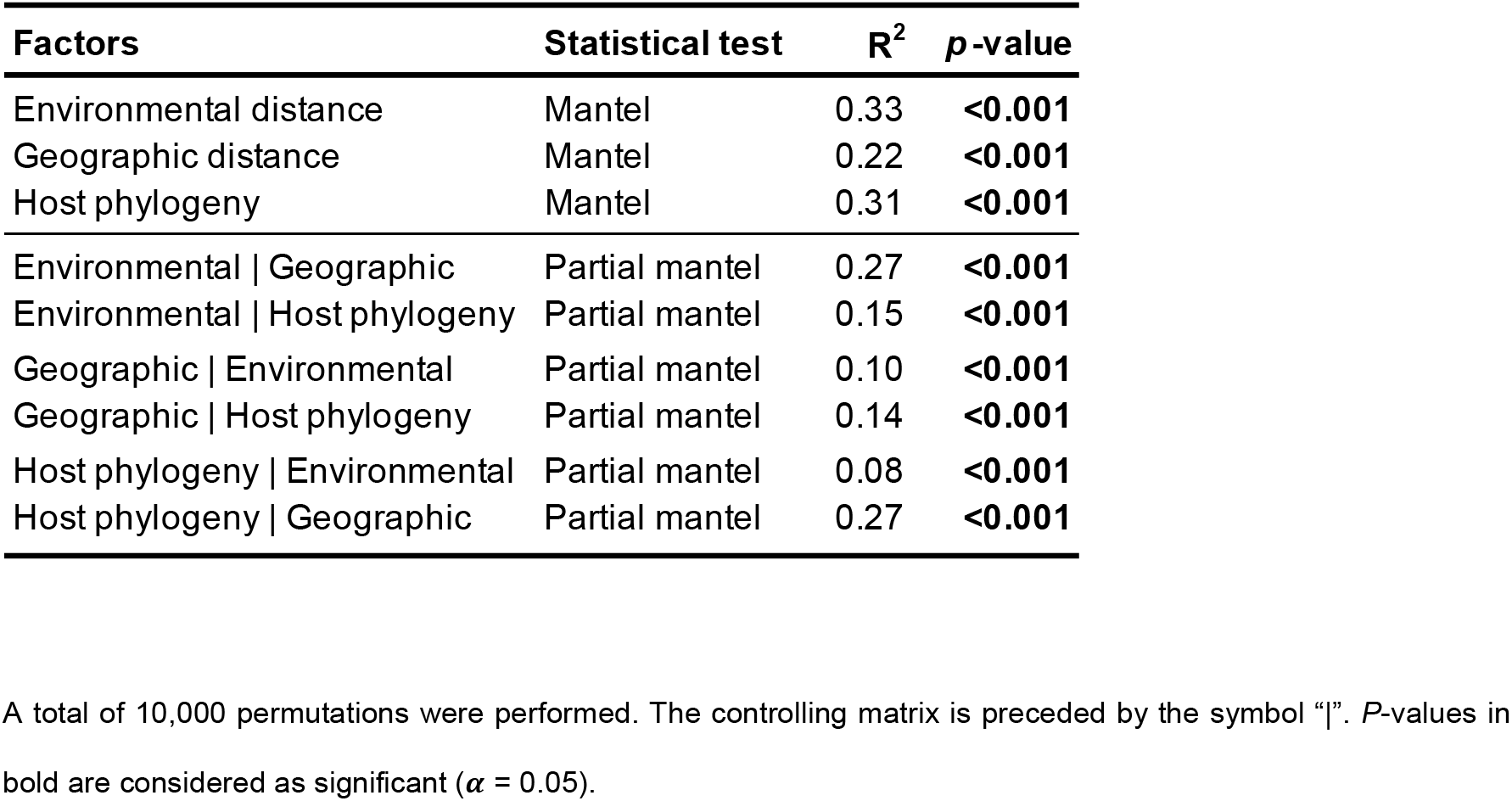
Mantel test analysis on GMB of Harpagifer spp.

### 3.4 Reduced core microbiome in *Harpagifer* spp. gut mucosa

A reduced core microbiome comprising 17 OTUs was detected across all *Harpagifer* species studied (Figure 3). This core microbiome represented in average 22.5% ± 2.9% of the relative abundance in gut mucosa samples, and spanned 9 bacterial classes belonging mostly to the *Gammaproteobacteria* phylum (Table 3). Prevalence of core OTUs ranged from 40% up to 96% of the gut mucosa samples. Two core OTUs were particularly abundant (OTU2 and OTU4, representing 12.6% and 6.9% of the total relative abundance in the whole GMM dataset, respectively), while the 15 others had relative abundance <0.9% (Table 3).

**Figure 3.**
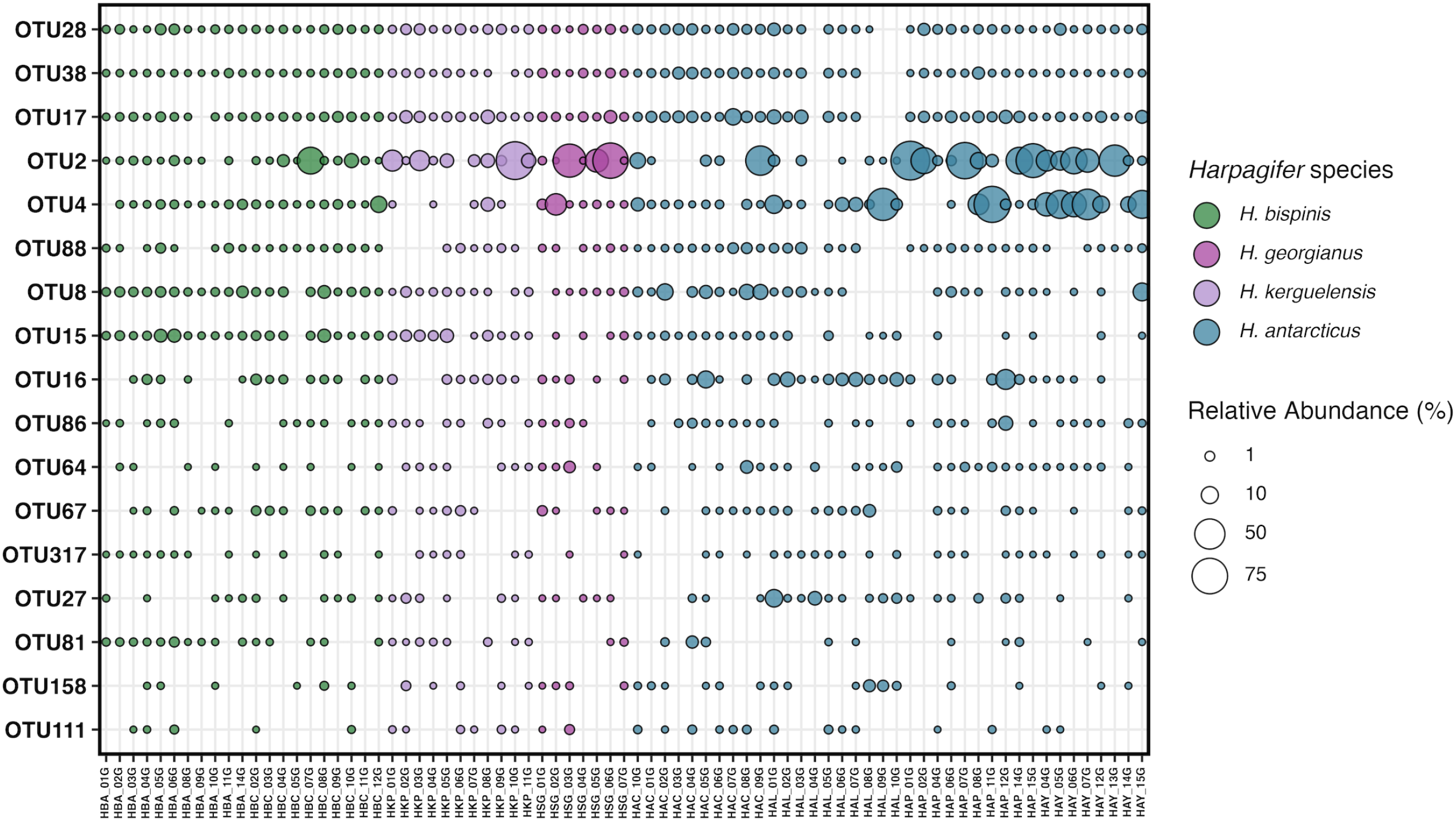
Bubble plot of core microbiome mucosa from *Harpagifer* sp. gut mucosa defined at >40% prevalence across samples. Relative abundances are given by sample. Colors are assigned to *Harpagifer* species. Details about taxonomic affiliation and relative abundance of these OTUs are provided in Table 3.

**Table 3.**
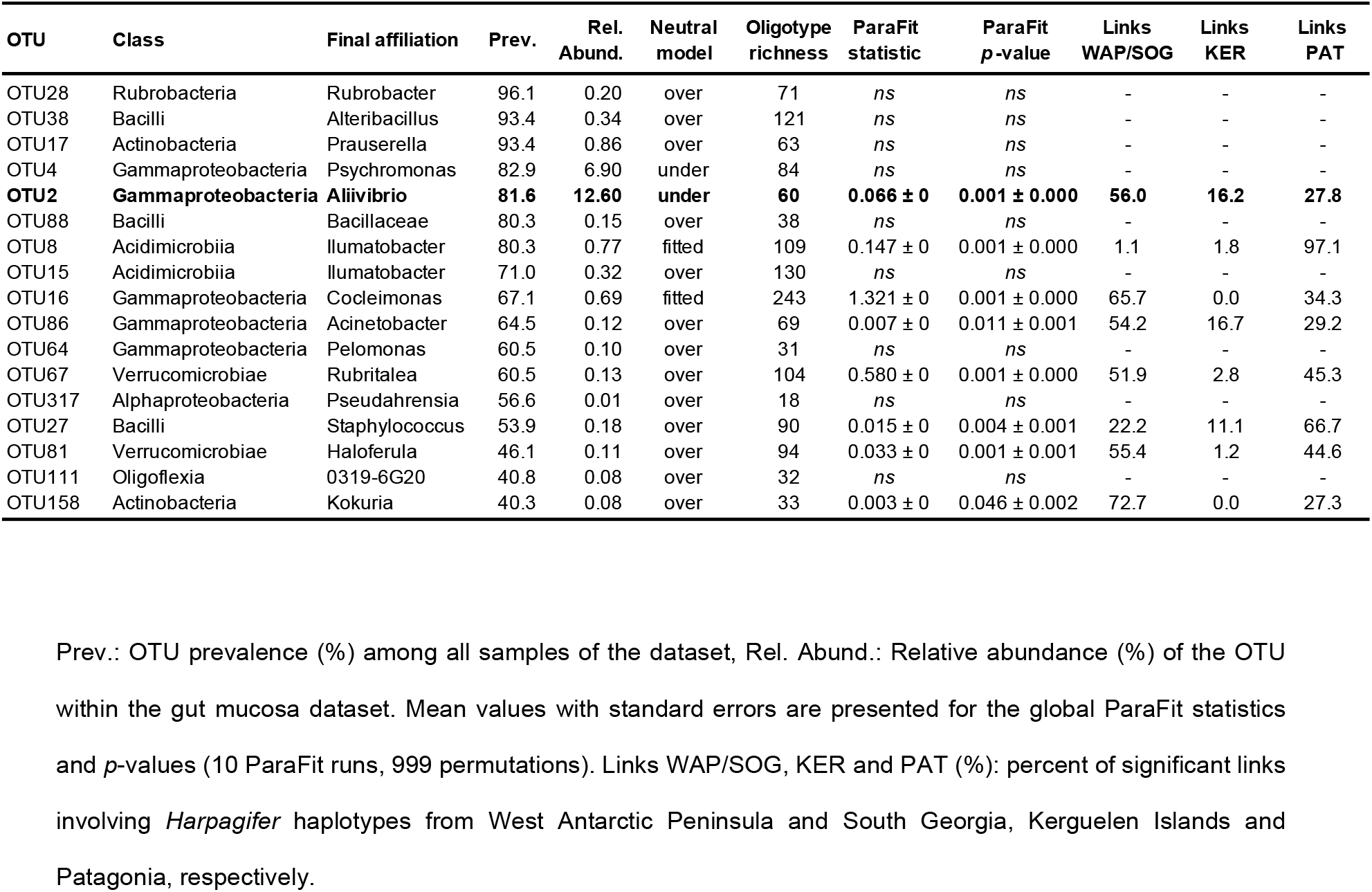
Fit of the core microbiome taxa from *Harpagifer* sp. gut mucosa to the co-phylogenetic model at the microdiversity level.

The fitting of GMM composition to the neutral model was relatively low, explaining no more than 20% of the gut microbiome variance of *Harpagifer* (*m=*0.004, R^2^=0.20, Figure 4). Remarkably, >88% of the core OTUs had an abundance-prevalence relationship that deviated from the neutral model predictions; most of them being either over- or under-represented (Figure 4 and Table 3).

**Figure 4.**
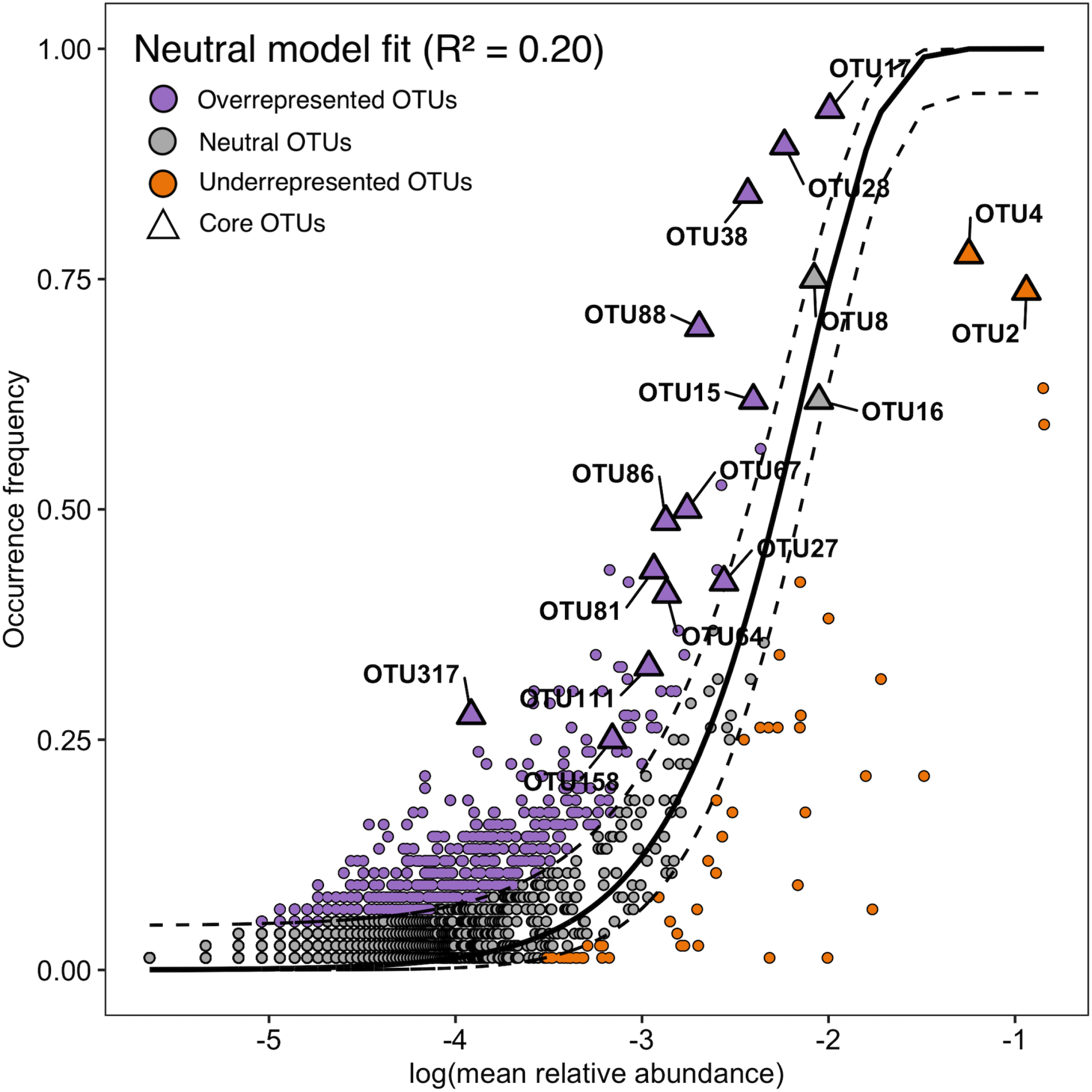
Neutral model fit for the gut mucosa microbiome (solid dark line) with bootstrap 95% CI (dashed dark line). OTUs in grey have a frequency of occurrence in the metacommunity congruent to their abundance under the neutral model hypothesis. OTUs that are above the 95% CI (purple) or below the 95% CI (orange) are significantly more frequent (overrepresented) or less frequent (underrepresented), respectively, than predicted by the model in the metacommunity of *Harpagifer* gut mucosa microbiome. Triangles represent the OTUs that were identified as part of the gut mucosa core microbiome.

The most abundant and prevalent OTU in the core microbiome of *Harpagifer* (*i.e.* OTU2), affiliated to *Aliivibrio*, occurred less frequently than the 95% confidence interval of the neutral model predictions, suggesting that this OTU is either selected against by the host (*i.e.* invasive taxa) or is especially constrained by dispersal limitation. Sequences affiliated to the *Aliivibrio* genus represented 51% of the GMM (393,084 sequences) and the OTU2 represented 99.4% of these sequences. The closest sequence retrieved from Blast analysis of the OTU2 representative sequence matched with an uncultured bacterium clone (99.3% identity), previously retrieved from the gut of *Notothenia coriiceps* (suborder Notothenioidei) fished in the Antarctic Peninsula (35).

### 3.5 Co-phylogeny between *Harpagifer* species and some core members of the gut mucosa microbiome

Eight out of the 17 core OTUs harbored significant signatures of co-phylogeny at the microdiversity level with *Harpagifer* spp. (*p*<0.05 for each OTU) (Table 3), suggesting that the evolution of these gut mucosa OTUs and their hosts was not independent (rejection of the H_0_ of Parafit test). The fit values were relatively low (ParaFit statistic<1.3), suggesting that external factors would also shape the observed patterns of co-phylogeny. Due to the absence of differences in GMM composition and host genetic divergence, WAP and SOG were combined in the results presentation of the host-oligotype links and co-phylogenetic representation. The proportion of significant links (*i.e.* associations between specific bacterial oligotypes and a given host species) involving WAP/SOG and PAT populations were globally similar, representing 47.4% ± 8.4% and 46.5% ± 8.6% of all significant links, respectively. Contrastingly, fewer links involving *Harpagifer* haplotypes from KER contributed to the global fit of the co-phylogeny model (8.3% ± 2.6%) (Table 3).

Due to its high abundance and prevalence in the GMM (deviating from neutral model predictions), and its significant co-phylogeny signal, a special interest was given to the *Aliivibrio* genus almost exclusively represented by the OTU2. The db-RDA analysis confirmed a significant influence of the host species identity on OTU2 oligotype composition (R^2^=0.07, *p*<0.01, Figure 5A). Pairwise PERMANOVA comparisons showed that the OTU2 oligotype composition was only different between *H. antarcticus* from WAP and *H. bispinis* from PAT (R^2^=0.05, *p*=0.001). The oligotype network of the OTU2 graphically confirmed that most of the predominant oligotypes were principally shared among SOG, WAP and KER regions, while the PAT oligotypes tended to be more exclusive to this region (Figure 5B).

**Figure 5.**
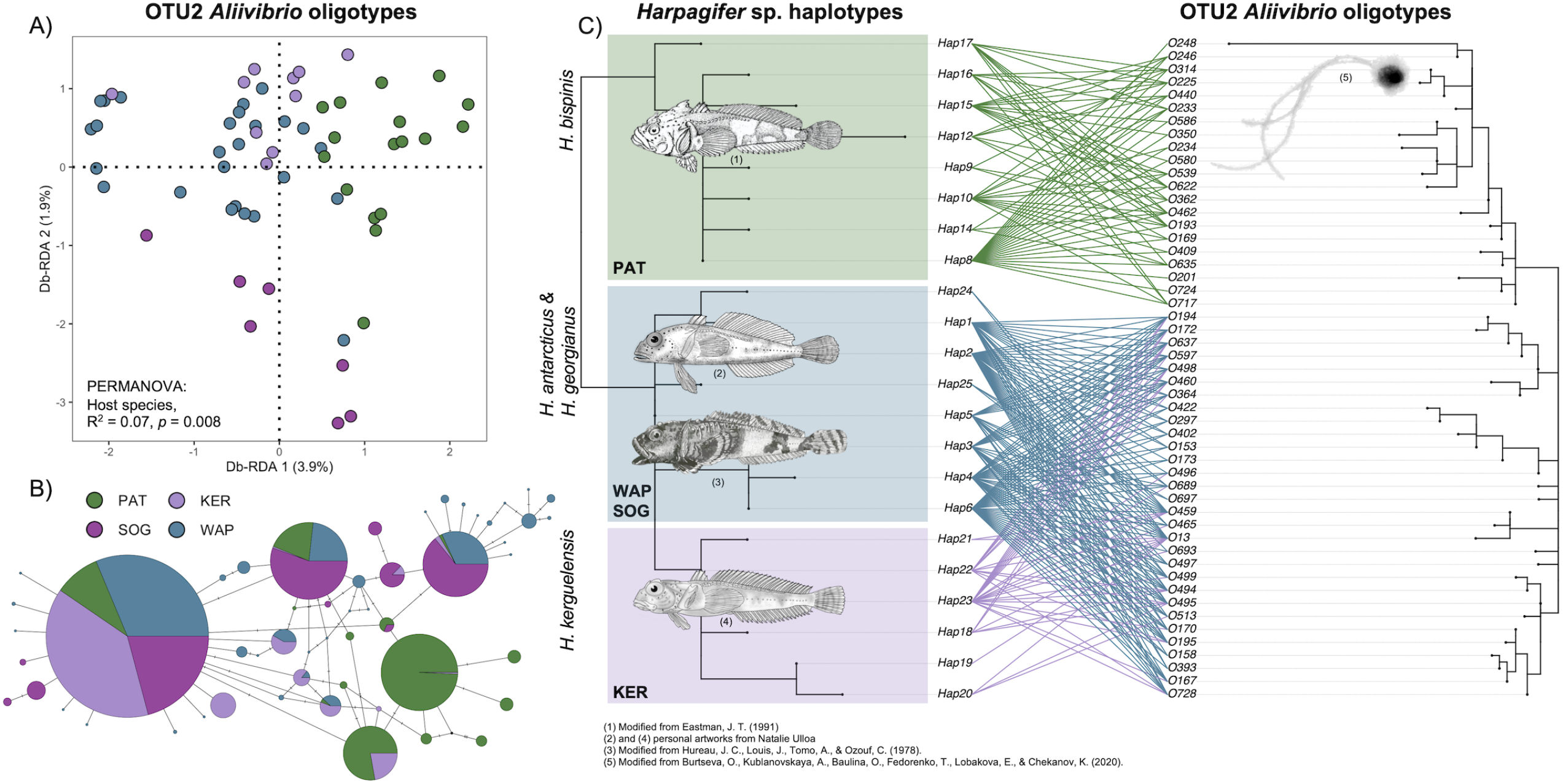
Example of co-phylogenetic pattern among *Aliivibrio* OTU2 oligotypes and *Harpagifer* spp. (A) Distance-based redundancy analysis (db-RDA) quantifying the contribution of the host species to explain the variations in OTU2 *Allivibrio* oligotypes composition in *Harpagifer* sp. gut microbiome. (B) Median-joining network of the OTU2 *Aliivibrio* oligotypes. Colors are assigned to the host biogeographic regions. The size of the circles is scaled on oligotypes’ frequencies. (C) Pruned tanglegram of the co-phylogenetic relationships between 22 *Harpagifer* haplotypes and 51 *Aliivibrio* oligotypes. Only the significant links between *Harpagifer* haplotypes *and Aliivibrio* oligotypes were plotted in the figure (ParaFit, p<0.05).

Beyond the qualitative effect of the host species identity, the phylogeny of OTU2 mirrors the phylogenetic patterns in *Harpagifer* (Figure 5C). The significant links involved 22 host haplotypes (out of the 25) and 51 *Aliivibrio* oligotypes (out of 60). The *Aliivibrio* oligotypes conformed two distinct clades, potentially representing below-genus divergence at the 16S V3-V4 locus. The *Aliivibrio* oligotypes significantly associated with *H. bispinis* clustered separately from the oligotypes significantly associated with all the other *Harpagifer* species, while *Aliivibrio* oligotypes significantly associated with *H. kerguelensis, H. antarcticus* and *H. georgianus* clustered together, consistently with the host haplotypes clustering (Figure 5C). PACo analysis of *Aliivibrio* oligotypes (OTU2) revealed that the ‘r2’ model led to the highest phylogenetic congruence between host and microbe phylogenies, suggesting that the *Aliivibrio* phylogeny is driven by *Harpagifer* phylogeny (and not the opposite), and that the degree of specialization of *Aliivibrio* oligotypes (quantified by the number of associations with *Harpagifer* haplotypes) also contributed to the global fit of the co-phylogeny model (Supplementary Figure 3).

## 4. Discussion

In this study, we investigated how the evolutionary changes among *Harpagifer* closely-related species associate with structural changes in their GMM. To the best of our knowledge, this study represents the first characterization at large-scale of the GMM of a fish genus inhabiting the SO. In natural systems, environment, geography and host phylogeny are generally confounded in the microbiota assembling among different species (5, 15, 71). Yet, controlling the environmental conditions to isolate the contribution of host genetics also distort the “wild” microbiome, since the host must be maintained in captivity (72, 73). Thus, combining a broad sampling strategy of *Harpagifer*, encompassing the three major biogeographic regions of its distribution area across the SO, and adapting statistical tools to explore the relative contributions of each factor (*i.e.* Mantel and MMRR), is a valid approach to provide a comprehensive evaluation of *Harpagifer* GMM assembly factors and to detect phylosymbiosis (74).

The *Harpagifer* host identity contributed to the variations observed in GMM composition, albeit to a low level. This result suggests that, despite their distribution in distinct biogeographic regions, the different *Harpagifer* species still share similar dietary and ecological constraints which lead to relatively similar gut microbial communities, as expected among species that have recently diverged in allopatry (31, 75). Beyond the mere host species identity effect, we showed that the phylogenetic distances among *Harpagifer* host species substantially correlate with gut bacterial dissimilarity distances, thus fulfilling the two central tenets associated with phylosymbiosis signature (15, 75). In other terms, the gut microbiomes were more similar among individuals of the same *Harpagifer* species than among different *Harpagifer* species (76). Numerous phylosymbiosis studies conducted among anciently diverged freshwater or marine fish species have led to the detection of weak and/or absent contribution of host evolutionary history on microbiome assembly (13, 30, 77-79). For instance, a relatively low phylosymbiosis signal was detected in skin microbiomes of 44 coral reef fishes encompassing 5 orders and 22 families (30), and very low and absent signals in the elasmobranchs and teleost subclasses, respectively (13). Contrastingly, by examining a recent radiation event in Central America lakes (0.5-1 Myr), Baldo et al. (80) deciphered a clearer signal of host genetics and phylogeography among the gut microbiome of 20 freshwater fish species from the monophyletic *Amphilopus* genus. In agreement with our hypothesis, the recent diversification of the *Harpagifer* genus across the SO is associated with a clear phylosymbiosis signal, thus expanding to marine fish the statement that phylosymbiosis strength among vertebrates and their microbiomes depends on the age of the last common ancestor (31). Although phylosymbiosis alone could arise from different evolutionary mechanisms unrelated to co-diversification, such as host filtering (81), its detection provides a strong basis for further investigating co-phylogeny of specific bacterial taxa shared among *Harpagifer* species.

Descriptions of core gut microbiomes in different species of wild marine fishes are rare and have been exclusively achieved so far across sympatric species from the same diet category (82–84). Huang *et al.* (85) did not find any common taxa in the gut microbiome of 20 marine fish species from coastal waters of Hong Kong, due to the high dependency of the gut core microbiome on the host’s feeding habits. Here, we revealed the existence of a core microbiome across gut mucosa of four *Harpagifer* species (77 individuals) distributed in three geographically-distant regions of the SO. This core was characterized by a relatively low diversity and the high dominance of a single taxon, as previously reported in both freshwater (77, 80) and marine fishes core microbiomes (85, 86). Even though we could not determine the persistence over time of the core taxa, the fact that most of these OTUs fell outside neutral model predictions suggests a major role of *Harpagifer* selective constraints in the recruitment, assembly and maintenance of its gut microbiome (80, 87).

Around half of the core bacterial taxa harbored strong signal of co-phylogeny with *Harpagifer* at microdiversity resolution. These taxa are expected to present a certain degree of host-specificity and to share at least some part of the host evolutionary history (5, 88). This result constitutes an unprecedented step forward in the understanding of marine fish holobiont, considering that most of the studies so far barely detected phylosymbiosis. Interestingly, the most abundant and prevalent taxon of the core GMM exhibited a significant co-phylogenetic pattern. This taxon belonged to the *Aliivibrio* genus, which has been identified by the past as a major component of the gut microbiome of notothenioid fishes (35, 37), and frequently associated with other fish species such as the European seabass and Atlantic salmon (89, 90). While there is no direct evidence of its ecological role in the fish holobiont, some authors suggest that *Aliivibrio* would be commensals, able to readily colonize the fish intestines (37, 91) and form biofilm onto intestine mucosa surface (92–94). This commensalism would be mediated by the capacity of *Aliivibrio* to degrade chitin, a highly conserved metabolism in the *Vibrionaceae* family, as confirmed by the detection of chitinase activity in *Aliivibrio* strains (95). Chitin is the most abundant biopolymer in the ocean (96), since it constitutes the exoskeleton of crustaceans such as copepods, amphipods and krill (97, 98), which are known to be part of the *Harpagifer* diet (47, 99, 100). Further, as the microbial metabolization of chitin aminopolysaccharide provides substantial source of carbon and nitrogen easily accessible for the host, a mutualistic cross-feeding is imaginable between *Harpagifer* and *Aliivibrio,* providing an ecological advantage to the holobiont (101, 102). Alternatively, other studies described the opportunistic and potentially pathogenic status of *Aliivibrio* strains inhabiting the intestinal tract of fishes (90, 103, 104). The deviation of *Aliivibrio* (*i.e.* occurring less frequently than expected by the neutral model analysis), suggests a selection against by the host, in line with a possible parasitic behavior of *Aliivibrio* (*i.e.* more abundant than expected) (65, 90, 105).

Since several *Aliivibrio* oligotypes co-occurred within the same *Harpagifer* species, and several *Harpagifer* species hosted the same *Aliivibrio* strain, we characterized the *Harpagifer*-*Aliivibrio* holobiont as a diffuse symbiosis (106). Unlike specialist symbionts that are highly specific to their hosts associations, more generalist symbionts generally show low phylogenetic congruence with their hosts (107). Accordingly, we found that the *Aliivibrio* oligotypes were less specific to the haplotypes of *H. kerguelensis* from KER, being frequently associated with *H. antarcticus* and *H. georgianus* haplotypes from WAP/SOG as evidenced by the links in the tanglegram representation. It is highly unlikely that this unspecific interactions’ pattern resulted from the unbalanced sampling of *Harpagifer* individuals across biogeographic region, as consistent numbers of host haplotypes were identified (i.e. WAP/SOG: 9, KER: 6, PAT: 10). We rather suggest that repeated host switch events generate the imperfect match between *Aliivibrio* and *Harpagifer* phylogenies. Although the transmission mode of *Aliivibrio* symbionts remains unknown, the diffuse symbiose and low phylogenetic congruence patterns suggest a horizontal transmission, either potentially widespread and repeatedly acquired from the surrounding environment, or vertically transmitted from parent to offspring (13), facilitating colonization of novel *Harpagifer* specimens and recurrent host-switch events. The frequent colonization of new individuals within a same fish species (*i.e*. *Salmo salar*) through the excretion of *Aliivibrio*-rich faeces has been previously proposed as an explanation to the wide distribution of *Aliivibrio* sp. in the surrounding waters (91, 103).

A strong co-phylogeny signal was observed between *Harpagifer* and *Aliivibrio*, mostly driven by the Patagonian and Antarctic/South Georgian oligotypes of *Aliivibrio* conforming two distinct sub-clades predominantly associated with their respective *Harpagifer* host species. According to PACo analyses, the most likely co-phylogenetic model was the adaptive tracking of *Harpagifer* phylogeny by *Aliivibrio*, suggesting that the diversification of *Aliivibrio* would result from unidirectional selection towards *Harpagifer* rather than independent response to a same biogeographic event or co-evolution (18, 69). Taking together, these results indicate that the biogeographic event experienced by *Harpagifer* (*i.e.* apparition of the Antarctic Polar Front (APF) (45)) leading to the speciation of *H. bispinis* by vicariance, indirectly generated the co-diversification of its specific *Aliivibrio* symbionts. An important prerequisite to this geographic model of co-diversification proposed by Groussin et al. (11) is the dispersal limitation of the microbial symbionts. The absence of significant links between *Aliivibrio* oligotypes from the PAT-related sub-clade with the *Harpagifer* haplotypes from WAP/SOG and KER, and the phylogeographic structure observed in *Aliivibrio* oligotypes network, support the relatively limited dispersal capacity between these two regions (65). Consistently, a previous study demonstrated that the APF limits the genetic connectivity among marine bacterial populations associated to invertebrate gut mucosa (*i.e.* sea urchin *Abatus*) from ANT and PAT (108). Contrastingly, some *Aliivibrio* oligotypes indifferently associate with *H. kerguelensis* and *H. antarctica*, further emphasizing that unlike host specimens, bacterial populations from ANT and KER may be occasionally interconnected generating a more diffuse pattern of association and counteracting *Aliivibrio* vicariance (108, 109). The synchronicity of *Harpagifer* and *Aliivibrio* divergence remains to be further explored using a robust time-calibrated phylogeny of the symbiont to confirm the strict co-diversification of hosts and symbionts speciation times (11). More generally, the absence of genetic and phylogeographic structures between *Harpagifer* populations of SOG and WAP regions, and the high homogeneity of *Aliivibrio* oligotypes’ composition attached to their gut mucosa, question the validity of the *H. antarcticus* and *H. georgianus* species. While further studies would be required to propose reunify H. antarcticus and H. georgianus species under a single Antarctic lineage, the contribution of gut microbiome and co-phylogenetic studies as an additional criteria to help in macroorganisms’ species and ecotypes delineation would represent a promising research avenue which may further promote the unification of the macro- and micro-biogeographic patterns examination (110, 111).

In conclusion, and contrastingly to the previously studied fish models, we revealed that host phylogeny was a substantial predictor of GMM composition of *Harpagifer*. Our survey represents the most conclusive evidence to date that phylosymbiosis and co-phylogeny occur between teleost fishes and their microbiome across the SO, and that recently diverged closely related species are suitable models to unravel phylogenetic congruency signals. We identified a small subset of bacterial taxa harboring robust co-phylogeny signal, largely dominated by *Aliivibrio*. These taxa are good candidates for further genomic-based exploration of their metabolic and ecological roles, due to their supposed tight and/or long-term interdependence with *Harpagifer*. While the co-diversification of *Harpagifer* and *Aliivibrio* remains to be confirmed, we provide a foundation to explore the mechanisms behind co-phylogeny signatures, notably by understanding the contribution of host biogeography into the diversification process of its symbionts.

## Declaration

### Ethics approval and consent to participate

Animal management was approved by the Universidad de Chile Institutional Animal Care and Use Committee (resolution n°20363-FCS-UCH).

### Consent for publication

Not applicable

### Availability of data and materials

Amplicon sequences of 16S rRNA and COI have been deposited in the National Center for Biotechnology Information (NCBI), under the Sequence Read Archive (SRA) PRJNA803378, and in GenBank under the accession number ON891147 to ON891171, respectively.

### Author contributions

EP, JO, and LC and GS conceived and developed the study. GS secured the research funds. EP, KG, TS, LC and GS contributed to the field work. GS performed the lab work and analyzed the molecular data. GS generated and analyzed the figures. GS drafted the manuscript and all authors revised it and approved its final version.

### Funding

ANID FONDECYT Postdoctoral Project (grant no. 3200036) ANID FONDECYT Regular Project (grant no. 1211672)

ANID – Millennium Science Initiative Program – ICN2021_002

## Acknowledgments

The authors thank the French Polar Institute project 1044 PROTEKER for field access, and the University of Wisconsin Biotechnology Center DNA Sequencing Facility (Research Resource Identifier – RRID:SCR_017759) for providing MiSeq® sequencing facilities and services. The bioinformatic analyses were partially supported by the infrastructure of the National Laboratory for High Performance Computing Chile (NLHPC). This research was partially supported by ANID Grant ACE210006. We would like to express our gratitude to Zambra Lopez, Francisco Bahamondes and Javier Cárcamo for their contributions in field sampling. We also thank Sebastián Rosenfeld and Natalie Ulloa for their graphical contributions to this study.

## Supplementary material

**Supplementary Figure 1. Redundancy analysis (RDA) ordination of the sampling localities based on a set of seawater properties collected in silico from the Bio-ORACLE database (v2.1).**

Dots are colored according to the sampling locality. The colored vectors indicate the contribution of each seawater variable on the two PCA dimensions. All these properties are mean values estimated at mean depth. Temp_range: Range of the seawater temperature, Temp_max: Maximum temperature, Temp_min: Minimum temperature, Temp_mean: Mean temperature, Chlo_mean: Chlorophyll A concentration, Carbonphyto_mean: mole concentration of phytoplankton expressed as carbon, pp_mean: net primary productivity of carbon, Nitrate_mean: mole concentration of nitrate, Salinity_mean: Salinity, Dissox_mean: mole concentration of dissolved molecular oxygen, Phosphate_mean: mole concentration of phosphate, Silicate_mean: mole concentration of silicate, Iron_mean: Mean mole concentration of dissolved iron.

**Supplementary Figure 2. Divergence of *Harpagifer* populations across sampled biogeographic regions.**

Median-joining haplotypes network computed from COI sequences of *Harpagifer* sp.

**Supplementary Figure 3. Phylogenetic congruence between *Harpagifer* hosts and OTU2 *Aliivibrio* according to the randomization algorithms implemented in PACo.**

The models tested were as follows: Host; host tracks the symbiont phylogeny, Symbiont; symbiont tracks the host phylogeny, Undetermined; unclear which phylogeny tracks the other, Specialist/generalist symbiont; symbiont tracks the host and the specialist/generalist feature of the symbionts also drive the co-phylogenetic signal.

**Supplementary Table 1. Abundance table of *Harpagifer* haplotypes across sampling localities.**

**Supplementary Table 2. Pairwise PERMANOVA on gut mucosa microbiome dissimilarities among Harpagifer species.**

F-statistics and *p*-values are provided on the lower and upper diagonal, respectively.

*P*-values are corrected with the Holm method.

